# Identification of a critical horseshoe-shaped region in the nsp5 (Mpro, 3CLpro) protease interdomain loop (IDL) of coronavirus mouse hepatitis virus (MHV)

**DOI:** 10.1101/2020.06.18.160671

**Authors:** Benjamin C. Nick, Mansi C. Pandya, Xiaotao Lu, Megan E. Franke, Sean M. Callahan, Emily F. Hasik, Sean T. Berthrong, Mark R. Denison, Christopher C. Stobart

## Abstract

Human coronaviruses are enveloped, positive-strand RNA viruses which cause respiratory diseases ranging in severity from the seasonal common cold to SARS and COVID-19. Of the 7 human coronaviruses discovered to date, 3 emergent and severe human coronavirus strains (SARS-CoV, MERS-CoV, and SARS-CoV-2) have recently jumped to humans in the last 20 years. The COVID-19 pandemic spawned by the emergence of SARS-CoV-2 in late 2019 has highlighted the importance for development of effective therapeutics to target emerging coronaviruses. Upon entry, the replicase genes of coronaviruses are translated and subsequently proteolytically processed by virus-encoded proteases. Of these proteases, nonstructural protein 5 (nsp5, Mpro, or 3CLpro), mediates the majority of these cleavages and remains a key drug target for therapeutic inhibitors. Efforts to develop nsp5 active-site inhibitors for human coronaviruses have thus far been unsuccessful, establishing the need for identification of other critical and conserved non-active-site regions of the protease. In this study, we describe the identification of an essential, conserved horseshoe-shaped region in the nsp5 interdomain loop (IDL) of mouse hepatitis virus (MHV), a common coronavirus replication model. Using site-directed mutagenesis and replication studies, we show that several residues comprising this horseshoe-shaped region either fail to tolerate mutagenesis or were associated with viral temperature-sensitivity. Structural modeling and sequence analysis of these sites in other coronaviruses, including all 7 human coronaviruses, suggests that the identified structure and sequence of this horseshoe regions is highly conserved and may represent a new, non-active-site regulatory region of the nsp5 (3CLpro) protease to target with coronavirus inhibitors.

**Importance:** In December 2019, a novel coronavirus (SARS-CoV-2) emerged in humans and triggered a pandemic which has to date resulted in over 8 million confirmed cases of COVID-19 across more than 180 countries and territories (June 2020). SARS-CoV-2 represents the third emergent coronavirus in the past 20 years and the future emergence of new coronaviruses in humans remains certain. Critically, there remains no vaccine nor established therapeutics to treat cases of COVID-19. The coronavirus nsp5 protease is a conserved and indispensable virus-encoded enzyme which remains a key target for therapeutic design. However, past attempts to target the active site of nsp5 with inhibitors have failed stressing the need to identify new conserved non-active-site targets for therapeutic development. This study describes the discovery of a novel conserved structural region of the nsp5 protease of coronavirus mouse hepatitis virus (MHV) which may provide a new target for coronavirus drug development.

## Introduction

Coronaviruses (CoVs) are enveloped, positive-strand RNA viruses which encode among the largest RNA virus genomes on the planet and infect a wide range of organisms including humans. To date, 7 human coronaviruses (HCoVs) have been identified, which include 4 HCoVs associated with seasonal common colds (HCoV-229E, HCoV-NL63, HCoV-OC43, and HCoV-HKU1) and 3 novel emerging coronaviruses associated with lower respiratory diseases and significant mortality (SARS-CoV, MERS-CoV, and SARS-CoV-2) (1). In December 2019, the first case of a novel coronavirus disease (COVID-19) was reported in Wuhan, China. Caused by a new emerging human coronavirus, now called SARS-CoV-2, this coronavirus likely emerged from bats and triggered a current worldwide pandemic resulting in over 8 million infections and over 400,000 deaths at the time of this writing (June 2020) (2). To date, there remains no commercially available vaccine for human coronaviruses and exhaustive efforts globally are underway to rapidly produce vaccines and effective therapeutic options to prevent and treat human coronavirus infections.

Coronaviruses encode positive-strand RNA (+ssRNA) genomes that range in size from 27 kb to 32 kb and represent among the largest RNA genomes (1, 3). During coronavirus infections, attachment and fusion of the virus are triggered by the viral Spike attachment protein (4–6). Upon entry and uncoating, the replicase open-reading frames of the virus, which encode up to 16 nonstructural proteins (nsps) are translated to form two variant polyproteins (pp1a and pp1ab) (**Fig. 1A**) (7–11). These polyproteins must undergo proteolytic cleavage by virus-encoded proteases, papain-like protease(s) (PLPs) and the nonstructural protein 5 (nsp5), to yield the mature replication machinery of the virus (1, 12–14).

**Figure 1:**
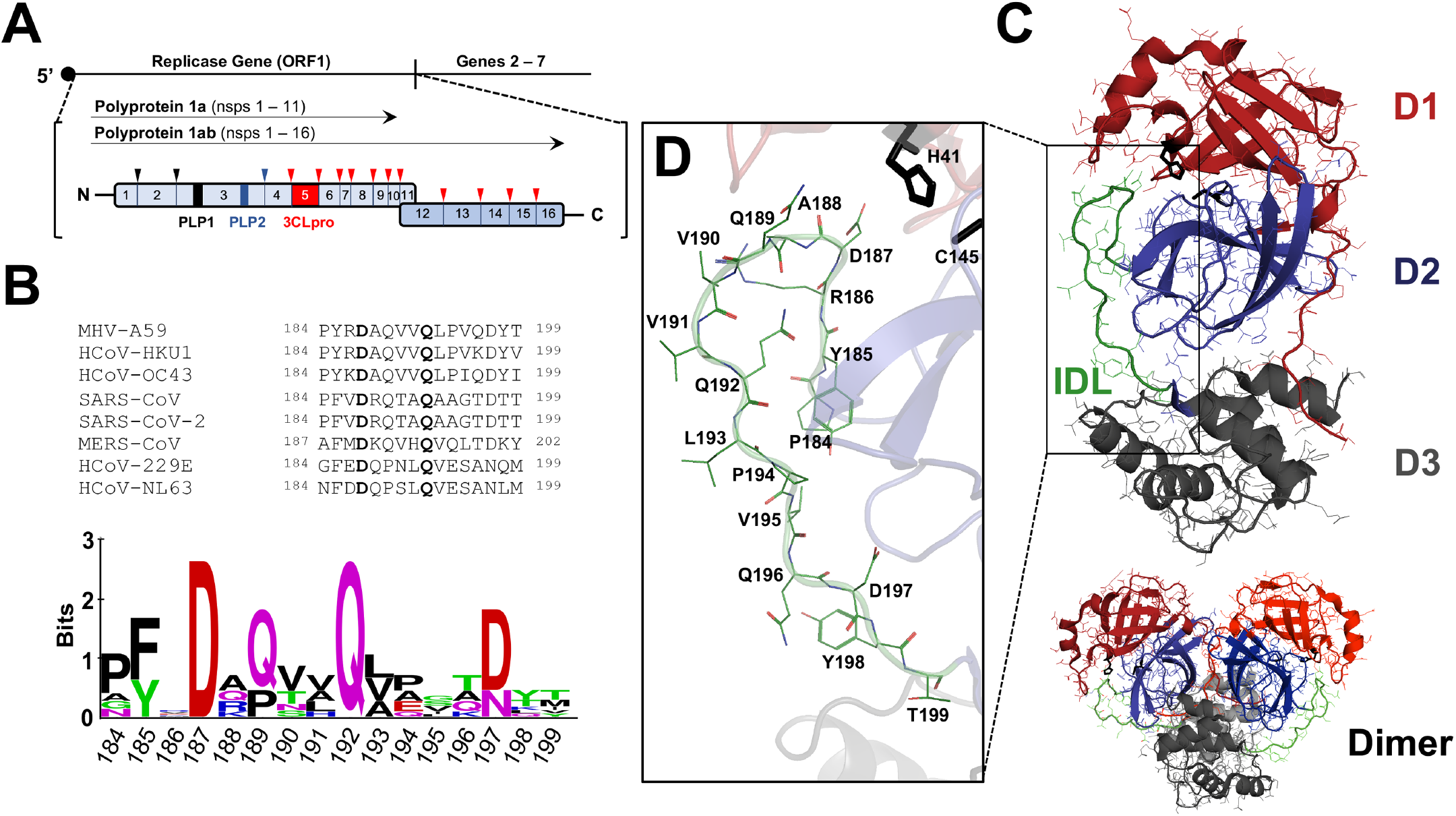
MHV nsp5 (3CLpro)-mediated polyprotein processing and conservation of structure and sequence of the nsp5 IDL. (**A**) Upon entry of the +ssRNA genome of MHV into cells, the first open-reading frame (ORF1) encoding the replicase machinery is translated into two variant polyproteins (pp1a and pp1ab) which undergo maturation cleavages by 3 viral encoded proteases (papain-like proteases 1 (PLP1, black) and 2 (PLP2, blue) and nsp5 (3CLPro, red)). The cleavage sites are marked by arrows with colors corresponding to the protease mediating each cleavage. (**B**) Sequence alignment (top) and sequence logo (bottom) of the MHV and 7 HCoV nsp5 IDL sequences. Residues D187 and Q192 are 100% conserved (in bold) in all known coronavirus nsp5 sequences identified to date. The sequence logo was generated using WebLogo with amino acid conservation represented in the height of each letter and the overall height of the amino acids represented in the position (31). (**C**) Crystal structure of an MHV nsp5 monomer and dimer (PDB – 6JIJ) with domains 1 (D1, red), 2 (D2, blue), 3 (D3, gray) and the D2-D3 IDL (green) color coded. (**D**) Expanded view of the MHV nsp5 IDL with the individual amino acids comprising the loop labeled along with the H41-C145 catalytic dyad residues.

Coronavirus protease nsp5 (M_pro_, 3CL_pro_) is a cysteine protease which is conserved in both overall structure and function among all coronaviruses identified to date (13, 15–17). Since its initial discovery, nsp5 has continued to be a primary target for design of coronavirus inhibitors and therapeutics. Nsp5 consists of three domains with domains 1 and 2 forming a chymotrypsin-like fold housing the His-Cys catalytic dyad and active site and a more divergent third domain promoting stabilization of the chymotrypsin-like fold and mediating nsp5 dimerization, an important event to complete 3CLpro maturation processing (**Fig. 1C**) (18–21). Connecting domains 2 and 3 is a 16 amino acid interdomain loop (IDL) which is a conserved feature in every nsp5 structure resolved to date (**Fig. 1B and 1D)**. Several recent studies using X-ray structures have suggested that residues located in the IDL form parts of the S2 - S4 substrate binding pockets for several human nsp5 proteases (22–25). Furthermore, biochemical modeling and docking of active site inhibitors to SARS-CoV, MERS-CoV, and the recent SARS-CoV-2 3CLpro proteases have suggested that engagement between residues found within the nsp5 IDL are critical for increasing affinity and inhibitory activity of active site inhibitors (24–26). Despite these biochemical studies and predictive models, there has yet to be any studies which have evaluated the function of the nsp5 IDL in a replicating virus. We hypothesize that the coronavirus nsp5 IDL represents an important structural and regulatory region of the protease.

In this study, we used site-directed mutagenesis to investigate how specific changes within the nsp5 IDL of mouse hepatitis virus (MHV), an established replication model for coronavirus study, impact overall virus replication (27). These studies provide the first detailed analysis of this conserved and important region of the nsp5 protease and may provide a new target for the development of coronavirus nsp5 inhibitors.

## Materials and Methods

### Cells, viruses, and antibodies

Recombinant wild-type (WT) mouse hepatitis virus (MHV) strain A59 (GenBank accession no. AY910861) was used as a wild-type virus control for the experiments described. Temperature-sensitive MHV nsp5 mutant virus S133A which was used as a temperature-sensitive control for efficiency of plating (EOP) analysis has been previously described (28, 29). Virus experiments were performed in murine delayed brain tumor 9 (DBT-9) cells, which are naturally permissive to MHV-A59 infection, and baby hamster kidney cells which express the MHV receptor (BHK-R) under selection with 0.8 mg/ml of G418. Complete Dulbecco’s Modified Eagle medium (DMEM, VWR) supplemented with 10% fetal bovine serum (FBS), 1% HEPES, and an antibiotic-antimycotic solution (Corning) containing penicillin, streptomycin, and amphotericin B, was used for growth of both DBT-9 and BHK-R cells. All biochemical experiments were carried out using rabbit polyclonal anti-nsp8 antisera previously described (29).

### Site-directed mutagenesis and recovery of MHV nsp5 IDL mutant viruses

The MHV infectious clone (MHVic) reverse genetics system used for the attempted recovery of the nsp5 IDL mutant viruses has been previously described by Yount *et al* (30). In brief, the nsp5 IDL mutations were engineered into the MHVic C fragment using a PCR-based approach with sense and antisense primers containing overlapping nucleotide changes corresponding to the desired amino acid changes in the nsp5 IDL. The MHVic C fragment sequences were all sequence confirmed prior to MHVic assembly, which involved the ligation of digested and gel purified cDNA fragments, *in vitro* transcription of the ligated cDNA (along with a N-gene containing cDNA) using an mMachine T7 transcription kit (Ambion), and subsequent electroporation into BHK-R cells. All virus recovery attempts were made at least 3 times and recovered viruses were expanded in DBT-9 cells and sequence confirmed before analysis.

### Viral replication assays and efficiency of plating (EOP) analysis

To evaluate viral replication kinetics, DBT-9 cells were grown to near confluency (~90 - 100%) in 6-well plates prior to infection with a virus multiplicity of infection (MOI) of 0.01 PFU/cell at either 37°C or 40°C. Throughout the replication time course, aliquots of virus were obtained and prewarmed media added back to ensure a constant volume. Virus titers were determined in duplicate by plaque assay on DBT-9 cells as previously described (29). Efficiency of plating (EOP) values were determined as the ratio of calculated titers by plaque assay at two different temperatures (40°C and 37°C) for the same aliquot of virus.

### Western blot analysis of nsp5 protease activity

DBT-9 cells were infected at an MOI of 0.5 with virus and cell lysates harvested in RIPA buffer (150 mM NaCl, 1% NP40, 0.5% Sodium deoxycholate, and 50 mM Tris pH 8.0) at 8 h p.i. Lysates were separated on a 4 - 15% polyacrylamide gel, transferred to a PVDF membrane, and blotted using an MHV nsp8-specific rabbit primary antibody, anti-nsp8 (VU123) (29). Western blots were resolved using an HRP-conjugated goat anti-rabbit secondary antibody and Western ECL substrate (Bio-Rad).

### Reversion analysis

A plaque assay was performed at 40°C in DBT-9 cells. After visible plaques had formed, 10 plaques were picked and were individually expanded in T25 flasks of confluent DBT-9 cells. At approximately 70 – 90% syncytial involvement, the viral RNA was isolated and used for sequencing of the nsp5 coding region.

### Sequence and structural analyses

The nsp5 IDL sequences of MHV-A59 and all 7 HCoVs were aligned using CLUSTALW and analyzed for sequence conservation using WebLogo (31). Structural analysis of coronavirus nsp5 proteases was performed using PyMol (The PyMOL Molecular Graphics System, Version 2.0 Schrödinger, LLC.) using the following protease structures available in the Protein Data Bank (PDB) or the Protein Modeling Data Base (PMDB): MHV-A59 (PDB 6JIJ), SARS-CoV (PDB 2Q6G), MERS-CoV (PDB 4YLU), SARS-CoV-2 (PDB 6M2N), HCoV-229E (PDB 2ZU2), HCoV-NL63 (PDB 3TLO), HCoV-HKU1 (PDB 3D23), and HCoV-OC43 (PMDB 0079872) (32–39).

### Statistical analyses

Differences in viral replication were evaluated by fitting the replication curves to logistic growth models. The replication curve data were log transformed and a three-parameter model was fit to each temperature condition by least squares (40, 41). Parameter estimates and 95% confidence intervals were calculated for each mutant strain. The parameters evaluated were maximum slope (replication rate), inflection point (time to maximal replication), and maximum titer. Strain parameter estimates with non-overlapping 95% confidence intervals were significantly different (p<0.05). EOP data were analyzed using one-way ANOVA with viral strain as the main effect. Post-hoc analysis was conducted using Tukey’s HSD to determine differences between strains. The data were log transformed to meet the assumptions of ANOVA but represented as non-transformed values for ease of interpretation. All statistical analysis were performed using JMP (SAS Institute, Cary, NC).

## Results

### Site-directed mutagenesis of the MHV nsp5 IDL reveals several residues and IDL modifications which fail to tolerate alanine-scanning mutagenesis

The MHV nsp5 interdomain loop (IDL) is comprised of 16 amino acids from P184 to T199. To assess the roles and contributions of the different residues and regions of the IDL to the nsp5 protease activity, we used a combination of alanine-scanning mutagenesis and C-terminal additions and deletions to initially mutate the MHV nsp5 IDL (**Table 1**). Of the 16 amino acids comprising the loop, a total of 8 virus mutants were successfully recovered (P184A, R186A, A188I, V190I, V191I, P194A, Q196A, and Y198A), 5 amino acid residues failed to permit virus recovery despite multiple attempts at rescue (Y185A, D187A, Q189A, Q192A, and T199A), and 3 amino acid residues were not evaluated (L193, V195, and D197). Among the unrecovered mutants, additional attempts to rescue using more conservative amino acid substitutions at residues D187 (D187E) and Q192 (Q192N) were also unsuccessful. A total of four different C-terminal modifications were also attempted, which included 2 different C-terminal additions (a duplication of residues 197 – 199 and a duplication of residue 199) and 2 different C-terminal deletions (a deletion of residues 197 - 199 and a deletion of residue 199). All four of these C-terminal modifications to the nsp5 IDL failed to permit virus recovery.

**Table 1:** Site-directed Mutagenesis of the MHV nsp5 IDL. Mutations were introduced and mutant virus recoveries attempted throughout the MHV nsp5 IDL using a reverse genetics system (30). Viruses which were recovered and those that did not permit recovery are described. Recovered viruses were sequence confirmed. Virus mutants which did not recover were attempted at least 3 times.

| MHV IDL Residue(s) | Virus Mutant Recovered | Virus Mutant Unrecovered |
| --- | --- | --- |
| **Pro184** | P184A |  |
| **Tyr185** |  | Y185A |
| **Arg186** | R186A |  |
| **Asp187** |  | D187A, D187E |
| **Ala188** | A188I |  |
| **Gln189** |  | Q189A |
| **Val190** | V190I |  |
| **Val191** | V191I |  |
| **Gln192** |  | Q192A, Q192N |
| **Leu193** |  |  |
| **Pro194** | P194A |  |
| **Val195** |  |  |
| **Gln196** | Q196A |  |
| **Asp197** |  |  |
| **Tyr198** | Y198A |  |
| **Thr199** |  | T199A |
| **C-term Insertions** |  | D197-T199 Ins, T199 Ins |
| **C-term Deletions** | | Δ197-199, Δ199 |

### Analyses of plaque formation, replication, and protease activity reveal a novel temperature-sensitive mutant in the MHV nsp5 IDL

To evaluate the replication kinetics of each of the recovered MHV nsp5 IDL mutants, we infected confluent DBT-9 cells with an MOI of 0.01 of each of the IDL mutants and titered aliquots over a 24 h period (**Fig. 2A**). All 8 recovered MHV IDL mutants exhibited indistinguishable replication kinetics compared to WT MHV. Previously, we described a total of 3 separate temperature-sensitive mutations (*ts*V148A, *ts*S133A, and *ts*F219L) in the MHV nsp5 protease whose phenotypes could be suppressed through long-distance second-site suppressor mutations (28, 29, 42). To evaluate whether any of the recovered MHV nsp5 IDL mutants may exhibit a temperature-sensitive phenotype, we performed an efficiency of plating (EOP) analysis by comparing the titers of each IDL virus by plaque assay determined at a physiologic (37°C) and elevated temperature (40°C) (**Fig. 3A**). Average EOP values were determined by the average ratios of titers at 40°C compared to 37°C, with those EOP values less than 10^-1^ indicating a greater than 10-fold reduction in titers at the elevated temperature as being temperature-sensitive (*ts)*. WT MHV exhibited an average EOP of 7.6 x 10^-1^. In contrast, previously described *ts* nsp5 mutant virus S133A, exhibited an average EOP of 1.49 x 10^-4^, consistent with the EOP previously reported (29). Two separate MHV nsp5 IDL mutants exhibited average EOP values less than 10^-1^ and were significantly lower than WT MHV (p<0.05): P184A and R186A. Mutant P184A exhibited an average EOP of 1.39 x 10^-2^. In contrast, IDL mutant R186A resulted in a much lower average EOP of 7.6 x 10^-4^, which was not significantly different from the known *ts* mutant S133A. No other IDL mutants exhibited average EOPs significantly different from WT MHV. These data suggested that mutagenesis of two separate IDL residues (P184A and R186A) have resulted in novel temperature-sensitive phenotypes. To determine whether the observed differences in phenotype for IDL mutants P184A and R186A are due specifically to defects in nsp5 protease activity or some other long-distance effect, we performed a Western blot to evaluate the ability for the P184A and R186A nsp5 proteases to process the maturation cleavage of a downstream replicase (pp1ab) protein, nsp8, during virus replication (**Fig. 3B**). Lysates from WT-, P184A-, and R186A-infected DBT-9 cells were compared for nsp5-mediated nsp8 processing at 37°C compared to 40°C. WT-MHV and P184A exhibited approximately equivalent levels (ratios of 1.08 and 0.99, respectively) of nsp8 protein detected at both temperatures. Consistent with its temperature-sensitive EOP, virus mutant R186A exhibited reduced nsp8 protein detected at 40°C compared to 37°C (ratio of 0.78) and when normalized to WT, exhibited an approximate 27% reduction in mature nsp8 protein produced at the elevated temperature. These data demonstrate that MHV nsp5 IDL mutation R186A is associated with reduced nsp5 activity at 40°C, whereas no appreciable difference in processing at 40°C was detected for mutant P184A.

**Figure 2:**
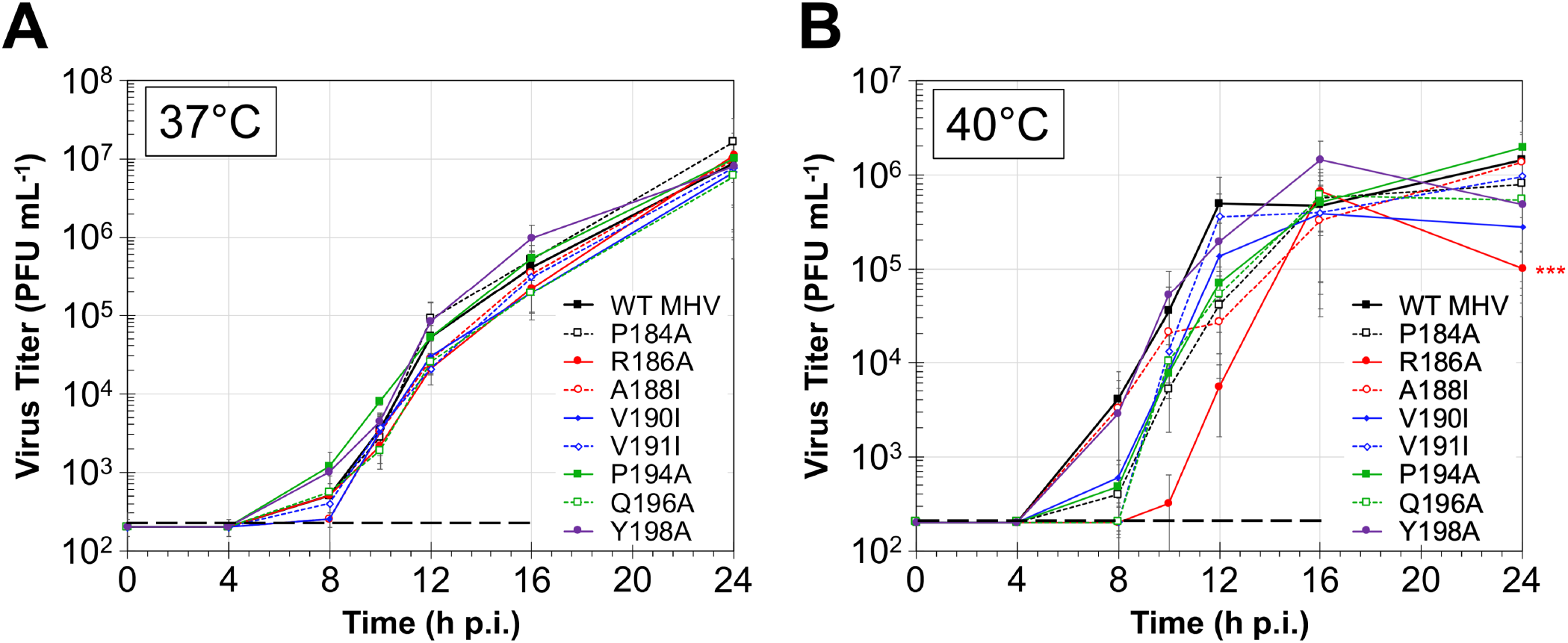
Replication analysis of MHV nsp5 IDL mutant viruses at 37°C (A) and 40°C (B). Confluent monolayers of DBT-9 cells were infected with an MOI = 0.01. Aliquots of viral supernatant were obtained over 24 h post-infection (h p.i.) and viral titers were determined by plaque assay in DBT-9 cells at 37°C. Data points in each graph represent the average titer ± SEM for 2 experimental replicates (**A**) or 5 experimental replicates (**B)**. The limit of detection is shown by a black dashed line and titers at or below the limit of detection were reported as equal to the limit of detection. Statistical analyses were performed to compare the time to reach the maximal replication rate with statistically significant times compared to WT indicated (***, p<0.05).

**Figure 3:**
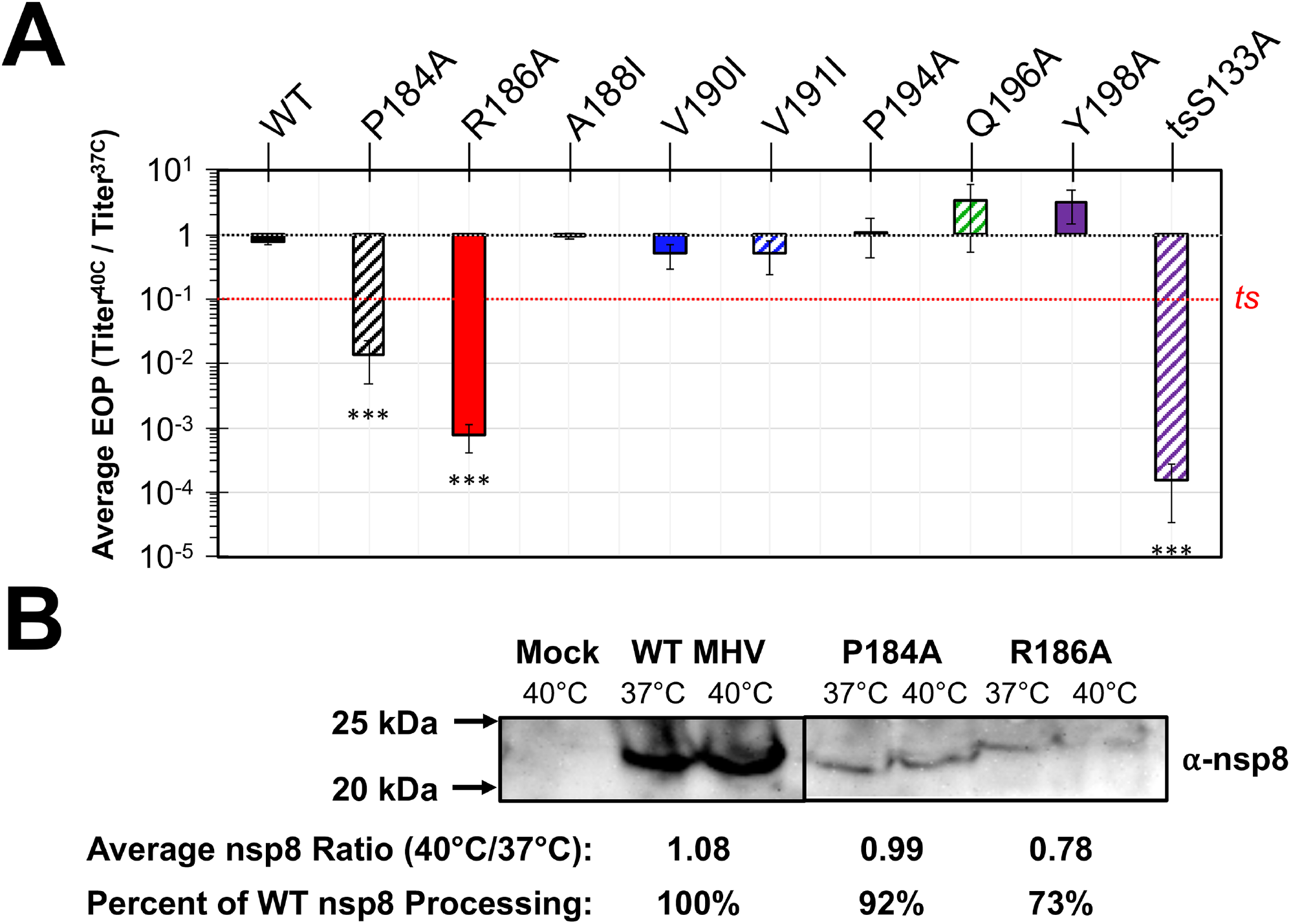
Efficiency of plating (EOP) and Western blot analysis of MHV nsp5 IDL mutant viruses. **(A)** Plaque assays were performed in confluent monolayers of DBT-9 to determine the capacity of each virus to form plaques at 37°C and 40°C. The efficiency of plating (EOP) was determined as the ratio of titers at 40°C to 37°C for an identical stock. The data shown reflect the average EOP ± SEM (N = 3). Significant differences from WT are indicated by asterisks (***, p<0.05, one-way ANOVA). **(B)** A Western blot was performed using mock or virus-infected lysates (WT, P184A, and R186A) which were harvested 8 h p.i. from DBT-9 cells infected at an MOI = 0.5. The protein lysates were resolved by SDS-PAGE, transferred to a PVDF membrane, and blotted for mature nsp8 (which undergoes maturation cleavage mediated by nsp5 protease) using anti-nsp8 antisera (VU123). Molecular weight markers are shown to the left of the bands. The average nsp8 band intensities (N = 2) were quantified using ImageJ and the ratios of nsp8 detected at 40°C to 37°C were reported for each virus as well as when normalized to the ratio for WT MHV. The two panels of the Western blot shown are from the same gel at the same exposure with irrelevant bands cropped out.

To assess the impact of elevated temperature on replication of the recovered MHV IDL mutant viruses, we repeated the MOI 0.01 replication assay in DBT-9 cells at 40°C (**Fig. 2B**). In contrast to replication at 37°C, the replication kinetics among the MHV IDL strains were far more variable, with most strains exhibiting a delay in logarithmic growth compared to WT MHV. Mutant P184A, which had shown a temperature-sensitive EOP of 1.39 x 10^-2^, failed to exhibit replication kinetics that were significantly different for wild-type or the other MHV IDL strains. In contrast, mutant strain R186A showed significantly delayed replication kinetics to reach the maximal logarithmic growth rate (p<0.05) compared to WT MHV consistent with its temperature-sensitive EOP of 7.6 x 10^-4^. Collectively, these data indicate that mutant R186A exhibits both significantly reduced capacity to form plaques and delayed replication kinetics at the elevated temperature of 40°C compared to WT MHV.

### Reversion analysis of *ts* MHV nsp5 IDL mutant R186A reveals three compensatory second-site suppressor mutations

To identify potential interacting residues and novel regulatory networks within the MHV nsp5 protease associated with residue R186, we performed reversion analysis at 40°C by expanding and sequencing formed plaques at the inhibitory temperature (**Fig. 4A**). A total of 10 plaques were selected at expanded in T25 flasks for virus collection and sequencing. Of these, 6 of these plaques resulted in the original R186A mutant virus while 3 of these plaques yielded R186A in addition to one of each of three different second-site putative suppressor mutations in nsp5: P184S, L141V, and L141I (**Fig. 4B**). Additional sequencing was performed on these 3 recovered viruses throughout the ORF1ab coding region and no other mutations were identified. The P184S mutation arose within the MHV nsp5 IDL, while residue L141 is located on the same loop housing the C145 catalytic residue of the active site.

**Figure 4:**
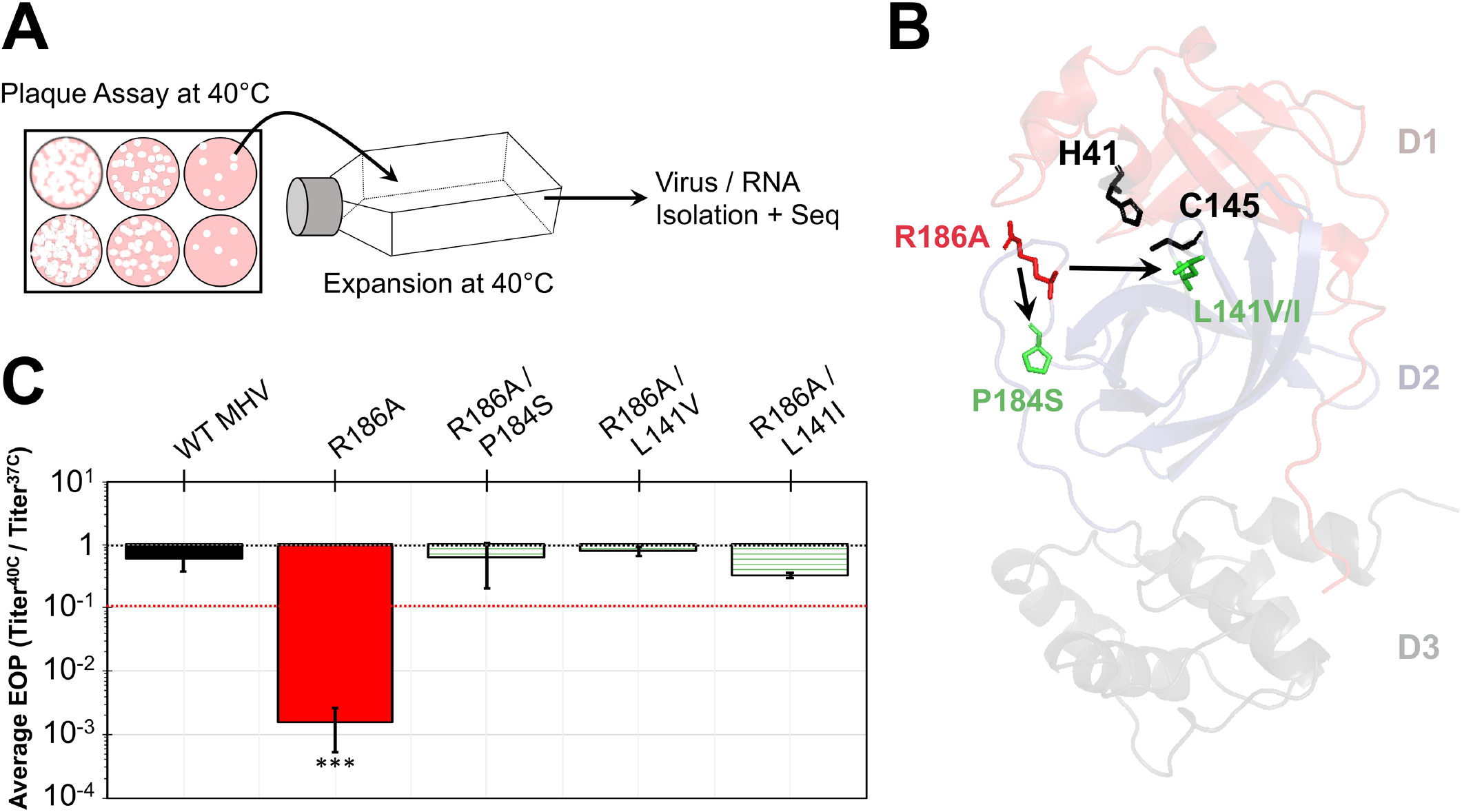
Reversion analysis of MHV nsp5 IDL mutant virus R186A. **(A)** Confluent monolayers of DBT-9 cells were infected with serially diluted mutant R186A for plaque formation at 40°C. A total of 10 plaques were picked and virus expanded at 40°C in T25 (25 cm^2^) flasks until approximately 30 – 50% involvement in syncytia. Viral RNA was isolated and sequencing performed throughout the ORF1ab replicase gene region. (**B**) Locations of recovered second-site revertant residues relative to the initial R186A mutation. (**C**) Plaque assays were performed in confluent monolayers of DBT-9 to determine the efficiency of plating (EOP) using titers at 40°C to 37°C for an identical stock. The data shown reflect the average EOP ± SEM (N = 2).

To evaluate whether the emergence of these second-site suppressor mutations aids in viral growth at 40°C, an EOP analysis was performed using these viruses at 37°C and 40°C (**Fig. 4C**). Consistent with earlier analysis, the R186A IDL mutant exhibited a temperature-sensitive and significantly reduced EOP (1.56 x 10^-3^) compared to WT MHV (0.60) (p<0.001). However, all 3 second-site suppressor mutant viruses (R186A/P184S, R186A/L141V, and R186A/L141I) resulted in indistinguishable EOP values (0.62, 0.78, and 0.33, respectively) from WT. These data collectively demonstrate that the addition of the second-site suppressor mutations was able to compensate for the initial defects in plaque formation associated with the primary R186A IDL mutation.

### The nsp5 IDL contains a structurally-conserved novel horseshoe region in the N-terminal region of the loop of human coronaviruses SARS-CoV, MERS-CoV, and SARS-CoV-2

To understand the structure and function of the IDL, we compared available crystal structures of human coronaviruses with MHV. To date, 6 of the 7 human coronavirus nsp5 proteases have been crystallized and resolved. We aligned these crystal structures along with MHV nsp5 and a modelled structure of HCoV-OC43 (**Fig. 5A**). Consistent with earlier studies, there was a high degree of conservation among domains 1 and 2 for all 8 of the proteases evaluated (most notably in and around the protein’s active site) (42). In contrast, domain 3 exhibited far more structural variability. The nsp5 IDL structure showed a high degree of structural similarity throughout including around a horseshoe shaped region in the N-terminus of the loop forming the inner and bottom part of the binding pocket for residues P2 – P5 of the substrate (**Fig. 5A and B**). Modeling using the crystal structure of SARS-CoV-2, residues D187 and T188 formed a distinct pocket in and around the P2 residue of Leu, residues T188 and Q189 establish the back wall of the P3 binding pocket, and residues Q189, T190, and Q192 are responsible for forming the back (Q189 and T190) and base (Q192) of the P4 and P5 binding pockets.

**Figure 5:**
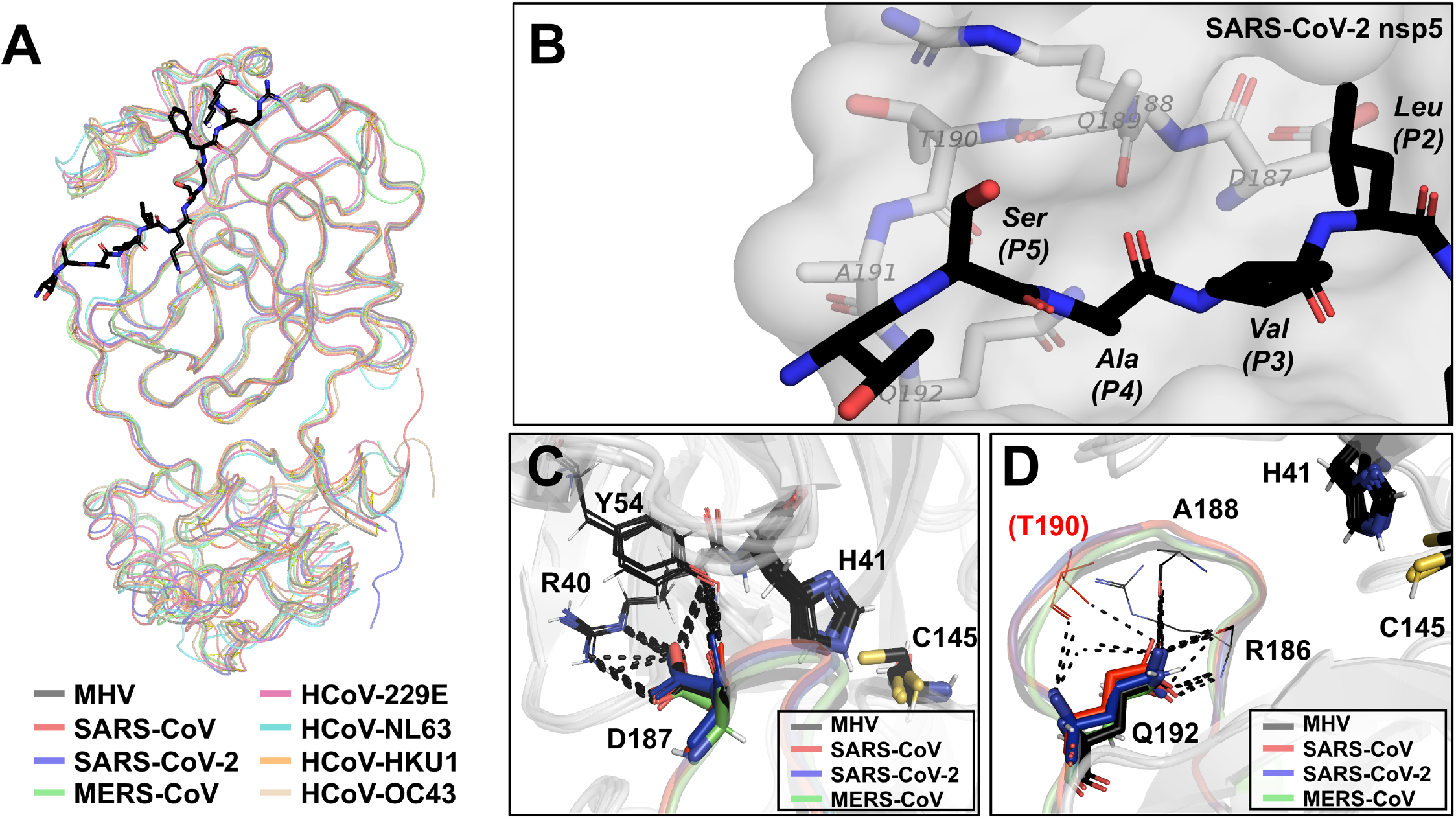
Structural analysis of the nsp5 IDL of human coronaviruses. **(A)** A ribbon overlay of nsp5 proteases of MHV (PDB 6JIJ) and the 7 HCoVs: SARS-CoV (PDB 2Q6G), MERS-CoV (PDB 4YLU), SARS-CoV-2 (PDB 6M2N), HCoV-229E (PDB – 2ZU2), HCoV-HKU1 (PDB – 3D23), HCoV-NL63 (PDB-3TLO), and the modeled structure of HCoV-OC43 (PM0079872) with a peptide encoding a SARS-CoV nsp5 autocleavage sequence (TSAVLQ↓SGFRKM) shown in black bound in the active site (32–39). (**B**) Modeling of the IDL-region of the substrate binding pocket with P2 – P5 residues labeled and the surface of SARS-CoV-2 nsp5 shown with contributing IDL residues D187 – Q192 identified. (**C and D**) Overlay of MHV, SARS-CoV, SARS-CoV-2, and MERS-CoV structures at the conserved MHV D187 (C) and Q192 (D) residues with the catalytic dyad H41 and C145 residues labeled. Predicted polar contacts between Q192 and other residues of the IDL are shown. SARS-CoV has an additional and unique predicted polar interaction with T190 (shown in red). All structures and modeling were performed with PyMol.

Among the MHV IDL mutants which failed to rescue were D187A, Q189A, and Q192A. Amino acid residues D187 and Q192 are structurally conserved in all sequenced nsp5 proteases to date (**Fig. 1B**). Both D187 and Q192 are located in a conserved horseshoe-shaped region in the N-terminus of the IDL. The D187 side chain projects from the top of the horseshoe-shaped region towards domain 1 and the protease active site and forms the inner wall pocket for the P2 binding site. In an alignment of the D187 residues of MHV, SARS-CoV, MERS-CoV, and SARS-CoV-2, the positioning and orientation of the side chain are highly conserved with predicted polar contacts with two additional highly conserved residues R40 (which is immediately adjacent to the catalytic H41) and Y54 (**Fig. 5C**). The Q192 side chain is conserved in its positioning towards the center of the horseshoe-shaped region where it shares predicted polar contacts with several other IDL residues including A188 and R186 (in MHV), R186 and R188 (in SARS-CoV-2), K191 (in MERS-CoV), and T190 (in SARS-CoV) (**Fig. 5D**).The coordination of Q192 with the backbone amino and carboxyl groups of R186 is conserved across all 4 viruses suggesting a potential role for the newly-identified temperature-sensitive residue. Collectively, these data indicate an important role of the horseshoe-shaped region of the IDL in forming the substrate binding pocket and stabilizing core of domains 1 and 2.

## Discussion

The coronavirus nsp5 protease remains a leading target for the development of inhibitor drug development. Over the last two decades, there have now been 3 emergent coronavirus outbreaks (including the current SARS-CoV-2 pandemic) which collectively highlight both the importance for rapid development of effective therapeutics for the treatment of COVID-19, but also the need to be prepared for potential future coronavirus outbreaks. In the present study, we evaluated the structure and function of the nsp5 protease IDL, a poorly studied and structurally conserved region of the protease. Using site-directed mutagenesis, we demonstrated that some residues and regions of the protease were capable of accepting mutations without apparent defects in viral replication, however a number of residues mostly located within a horseshoe-shaped region in the N-terminus of the protease either failed to permit virus recovery or resulted in a viral temperature-sensitivity. Of the 16 amino acid residues comprising the loop, we were able to successfully recover viral mutants at 8 different locations (**Table 1**).

Despite the overall structural conservation of the entirety of the loop, the majority of these mutations resulted in no apparent defects in viral replication compared to WT. A few of these residues (A188, V190, and V191) with no apparent viral defects are known to form the basis of part of the P3 - P5 substrate binding pockets of the protease (24, 26). Yet, compared to the rest of the IDL, these residue positions showed among the least sequence conservation (**Figure 1B**), which may explain the plasticity with which these residues could tolerate mutagenesis as well as cleavage site variability among coronaviruses (16). Similarly, more C-terminal residues P194, Q196, and Y198 are also found in more variable sequence locations within the IDL. Collectively, these 8 residue positions may simply represent flexible linker residues than serving additional structural supportive or enzymatic roles within the protease.

Residues P184 and R186, while rescued when mutated to alanine amino acids, exhibited reduced capacity to form plaques at 40°C. P184 is found at a bend leading into the horseshoe shaped region of the IDL and may be responsible for helping stabilize the N-terminal anchor of the loop within domain 2. Replication analysis and Western blots of the P184A mutant virus failed to show significant differences from WT MHV, however the selection of a P184S mutation in reversion analysis of R186A may suggest that these two residues represent stabilizing and interacting nodes within the protease (**Figure 4B**). We previously described 3 different temperature-sensitive mutations in MHV-A59 (S133A, V148A, and F219L) which all shared overlapping compensatory second-site suppressor mutations (28, 29, 42). All 3 viruses selected for an H134Y mutation, while the temperature-sensitive V148A mutation selected for an S133N mutation. Furthermore, second-site mutations were identified for F219L which were located greater than 20 Å away from the initial mutation. P184A is located on an adjacent loop in domain 2 to both S133 and H134 (less than 6 Å) in distance (not shown). MHV viral mutant R186A was found to exhibit delayed replication kinetics (**Figure 2**), reduced capacity to form plaques (**Figure 3**), and reduced nsp5-mediated proteolytic processing at the elevated temperature of 40°C, consistent with a temperature-sensitive phenotype (**Figure 3**). Perhaps surprising, the R186 residue position was the most variable and least conserved structurally among all 7 HCoVs evaluated (**Figure 1B**). Structural analysis of the MHV, SARS-CoV, SARS-CoV-2, and MERS-CoV revealed that the side chain of the 100% conserved Q192 appears to form conserved polar interactions with the backbone amino and carboxyl termini of the residue 186 position (**Figure 5D**). These data may suggest that Q192 is stabilized within the horseshoe shaped region of the IDL through anchoring to the residue 186 position immediately across from it. In addition, the selection of a compensatory change in the P184 position during reversion analysis would support a role of both the 184 and 186 residue positions in mediating loop stabilization. In addition to a P184S second-site suppressor mutation, two additional mutations (L141V and L141I) were also each associated with compensating for the R186A temperature-sensitivity (**Figure 4B**). Residue L141 is part of a well-characterized 3-residue loop (S139 – L141) that forms part of the S1 subsite of the protease and transitions into a short 3_10_-helix triggering an inactive conformation that has been shown in SARS-CoV serving as a putative enzymatic switch from inactive to active conformations. Introducing a L141T mutation into an inactive SARS-CoV triple mutant (G11A/R298A/Q299A) resulting restoration of nsp5 protease activity (20, 43). These earlier studies suggest that L141, helps regulate substrate accessibility to the active site. I141, one of the residues selected for by R186A in MHV, is the wild-type residue in two different a-CoVs: HCoV-229E and HCoV-NL63. The selection for a mutation at this residue position to restore full activity of R186A at 40°C may indicate that the protease experienced instability in or around the active site and substrate binding pocket of the protease.

There were 5 IDL residues which failed to permit MHV mutant virus rescue (Y185A, D187A/E, Q189A, Q192A/N, and T199A) (**Table 1**). Residue T199, while far more variable, is located at the C-terminal end of the IDL and efforts to modify this region by either additions or deletions were not tolerated. These data may suggest an important role of T199 in stabilizing the base and positioning of the IDL. Comparatively, four of these residue positions (185, 187, 189, and 192) show a high level of amino acid conservation with two of these residues (D187 and Q192) being 100% conserved across all known coronavirus nsp5 protease sequences to date (**Figure 1B**). All four of these residues are found within a conserved horseshoe-shaped region within the N-terminus of the nsp5 IDL. We propose that this horseshoe-shaped region is a critical region of the protease for both structure and function based on the following observations: (1) A total of 6 different residues either failed to tolerate mutagenesis (Y185, D187, Q189, and Q192) or resulted in altered phenotypes under elevated temperatures (P184 and R186); (2) Biochemical analysis and predictive structural modeling indicates that multiple residues including D187 – Q192 are involved in forming the P2 – P5 substrate binding pockets; (3) Predicted polar interactions shown here between D187 and conserved domain 1 residues R40 and Y54 appear to stabilize the positioning of the loop which is conserved in all nsp5 protease structures to date; and (4) Reversion analysis of temperature-sensitive IDL mutant R186A selected for two separate mutations at residue L141, which is known to regulate substrate accessibility to the active site and enzymatic activation. Previous biochemical analyses have suggested that next-generation coronavirus nsp5 inhibitors need to coordinate with the P2 – P5 substrate binding site to increase affinity and efficacy. The IDL data presented here support this idea and highlights a novel, conserved region found in all 7 HCoVs to date, including the recent emerging and pandemic SARS-CoV-2.

## Acknowledgements

This work was supported by a grant from the Holcomb Awards Committee (HAC) of Butler University (C.C.S.) and funding and support from the Butler University Department of Biological Sciences (C.C.S.). We would like to especially thank Dia Beachboard, Lindsay Maxwell, and the other members of the Stobart and Denison Labs for their support and guidance throughout this multiyear endeavor.

